# Preliminary findings of single session of noninvansive brain stimulation over parietal lobe and performance on spatial memory task

**DOI:** 10.1101/736892

**Authors:** V Marija Čolić, Uroš Konstantinović, Jovana Bjekić, R Saša Filipović

## Abstract

Spatial memory relies on efficient encoding, storage and retrieval of spatial information, which enables us to remember paths or locations of objects in everyday life. Moreover, this type of memory has been shown to decline with age and various neurodegenerative disorders. Parietal cortex has been shown to play an important role in the formation of short-term representations of spatial information. The aim of the current study was to test the possibility of immediate and long-term spatial memory enhancement, by increasing excitability of parietal posterior cortex. We used transcranial direct current stimulation (tDCS) over posterior parietal cortex in a placebo-controlled cross-over study. Participants received anodal (1.5 mA) or sham tDCS stimulation over P4 site (10-20 EEG system) for 20 minutes in two separate sessions. Immediately after stimulation, participants completed a spatial maze task, which consisted of learning block, 2D recall, and 3D recall. Spatial memory performance was tested 24 hours and 7 days after stimulation, to assess potential long-term effects. We found no significant effects of anodal stimulation on spatial memory performance either on immediate or delayed recall. This was the case with both, 2D and 3D spatial memory recall. Our results are in line with some studies that suggest that single brain stimulation sessions do not always produce effects on cognitive functions.

## Introduction

Spatial memory involves the encoding, storage, and retrieval of spatial information, which is crucial for everyday life functioning (Allen, 2004). Efficient spatial memory use enables us to remember paths, locate objects in our environment, use a maps, etc. Spatial memory is not unitary and it includes many specific cognitive mechanisms for processing different aspects of spatial information e.g. object recognition, spatial orientation, spatial navigation, object location binding, etc. It is important to distinct between memory for spatial layouts, such as involved in object location memory, and sequential processing of spatial information like a path learning (Schachter & Nadel, 1991). In terms of spatial representations there is a traditionally accepted distinction between allocentric and egocentric spatial frames. During allocentric framing, objects and their locations are being processed in reference to other objects, while egocentric framing includes self-referential processing of spatial information (Burgess, 2006). It is also relevant to differentiate spatial memory from spatial navigation which represents a complex ability to find and maintain a route from one place to another (Allen, 2004).

Neural basis of spatial cognition is highly investigated and there are two brain regions consistently showing substantial role in spatial information processing. The first one is hippocampus which plays a crucial role in all memory processes. Lesion studies showed crucial contribution of hippocampal activity for processes like path learning, accuracy metric in allocentric framing, and object location encoding (Kessels, De Haan, Kappelle, & Postma, 2001). Several theories have been linking hippocampus with spatial memory, assuming hippocampal function as allocentric cognitive map (O’Keefe & Nadel, 1978) or a binding device (Eichenbaum & Bunsey, 1995) integrating different contextual features of information in the environment. The other neural structure involved in spatial memory processes is posterior parietal cortex (PPC). It has been shown that PPC has crucial role in generation of short-term representations of spatial information (LaBar, Gitelman, Todd B. Parrish, & Mesulam, 1999; Roy, Sparing, Fink, & Hesse, 2015). PPC is also taking important part in egocentric framing (Bird & Burgess, 2008; Byrne, Becker, & Burgess, 2007), as it has function of integrating multiple sensory information (Andersen, 1997).

Additionally, it is traditionally suggested that spatial memory functions are strongly lateralized, and that right hemisphere has a key role in spatial processing. Namely, there is evidence that right hippocampus is involved in spatial locations recall (Milner, Johnsrude, Crane, Trans, & Lond, 1997; Smith & Milner, 1989) as in allocentric and egocentric framing in spatial navigation through computerized environment (Antonova et al., 2009). On the other hand, empirical findings about PPC lateralization in spatial memory functions (Baciu et al., 1999; Kessels, Kappelle, Haan, & Postma, 2002; van der Ham, Raemaekers, van Wezel, Oleksiak, & Postma, 2009) reveals bilateral PPC activity, considering two types of spatial relations processing. Although the data are not always consistent it seems that categorical spatial relations are processed mostly in left hemisphere, while right hemisphere seems to be more dominant in coordinate spatial relations processing.

In last two decades, development of non-invasive brain stimulation techniques has enabled targeted enhancement of different cognitive and motor functions in both healthy and clinical populations. The main advantage of this methodology is the possibility to safely (Antal et al., 2017; Grossman, Woods, Knotkova, & Marom, 2019) conduct controllable experiments which can provide causal evidence about neural basis of cognitive functions, unlike correlational neuroimaging techniques and less-controllable lesion studies. In this study, we used transcranial direct current stimulation (tDCS), which is widely used in scientific and clinical purposes (Berryhill & Martin, 2018). This procedure assumes generation of electrical field over the targeted brain area which changes ions distribution of nerve cells in targeted brain region and gives an outcome of few millivolts change of the neuronal resting potential (Stagg & Nitsche, 2011). Thus, application of tDCS gives possibility of increasing cortical excitability of the neurons in specified brain area which leads to higher probability of neural “firing” of targeted region (Nitsche et al., 2008). Anodal tDCS tends to induce increase of excitability of the underlying neural tissue (Nitsche & Paulus, 2000), which is usually linked with better performance on cognitive measures (Brunoni & Vanderhasselt, 2014; Summers, Kang, & Cauraugh, 2016) It is assumed that physiological mechanisms of tDCS effects lay in modulation of synaptic plasticity of GABA and glutamate systems.

The aim of the study was to investigate the effects of parietal tDCS on spatial memory performance. We tested whether the single-session anodal tDCS over right PPC would enhance spatial memory performance immediately after the stimulation. Furthermore, we performed assessments 24 hours and 7 days later to test for potential long-lasting after-effects.

## Methods

### Participants

The sample consisted of twenty-two healthy right-handed University students (12 female and 10 male, aged 20-28). Participants were preselected in line with tDCS inclusion criteria (Antal et al., 2017) and all of them were native speakers, with normal or corrected-to-normal vision. None of them reported previous history of head trauma, neurologic or psychiatric disorders. Study was approved by the Institutional ethics board and all participants gave their written informed consent.

### Transcranial direct current stimulation

The tDCS was delivered by STMISOLA (BIOPAC Systems, Inc., USA), controlled by CED1401 Plus using Signal software (Cambridge Electronic Design, Cambridge, UK), via rubber electrodes (5×5 cm) encased in a pair of saline-soaked electrode sponge pockets. To modulate PPC activity, anode was placed over P4 site of the International 10-20 system of EEG electrode placement, while reference electrode was placed over the contralateral cheek. P4 has been commonly used for targeting right PPC in different tDCS studies (Ghanavati, Nejati, & Salehinejad, 2018; Roy et al., 2015; Wright & Krekelberg, 2014). In the active condition, the 1.5mA constant current was delivered for 20 minutes, with a gradual ramp up/down over first and last 60 seconds, respectively. This procedure conformed to contemporary safety standards and produces no significant adverse effects as shown in multiple studies (e.g. Brunoni et al. 2011; Matsumoto and Ugawa 2017; Nitsche & Paulus, 2011). The sham condition followed the same routine except for the current being administered for only 60 seconds at the beginning and at the end of the treatment (gradual ramp-up/down), making it indistinguishable from the real stimulation (Nitsche et al., 2008). The order of stimulation conditions was counterbalanced across participants (i.e. half of participants first received anodal stimulation, while the other half first received sham).

### Spatial memory task

To assess spatial memory performance, we constructed two parallel forms of spatial maze task which was consisted of learning block, two-dimensional recall block (2D) and three-dimensional recall block (3D). In the learning block, participants were instructed to memorize three sequentially presented paths on a 7×7 grid, resembling street map. The paths were presented on the computer screen separately and gradually step-by-step, from starting to the final point, with a one second per step rate. Step was defined as a part of the path which lay between two “crossroads” on the grid. Three paths differed on color (green, purple and blue) and number of the steps (10, 6, 14 respectively). Both forms used the same coloring and number of the steps for each path, but differed regarding to starting and final points. After the third path had been presented, all three paths were simultaneously shown on the screen for three seconds (Figure 1). Before continuing with the next block, a visual noise picture was presented on the screen for two seconds. In the beginning of the 2D recall block, participant was shown a starting point on the grid and instructed to reproduce (step by step) each path by pressing arrow keys on the keyboard. One step of the path or the red letter “X” would appear as a feedback after every correct or incorrect response, respectively. In the 3D recall block participant was presented a computerized 3D maze environment and a color cued starting point to indicate which path was required to reproduce (Figure 2). Participant was moving through the maze by pressing arrow keys on the keyboard. At the moment when participant faced the “crossroad”, red question mark would appear on the screen, forcing him to choose one of three sides of the path. If responded correctly, participant would continue to move through the maze. Conversely, red “X” would appear as a mistake feedback. Final 2D and 3D scores were calculated as a sum of all correct responses that were given without mistakes for all three paths. The highest possible raw score for each condition was 30.

**Figure 1.**
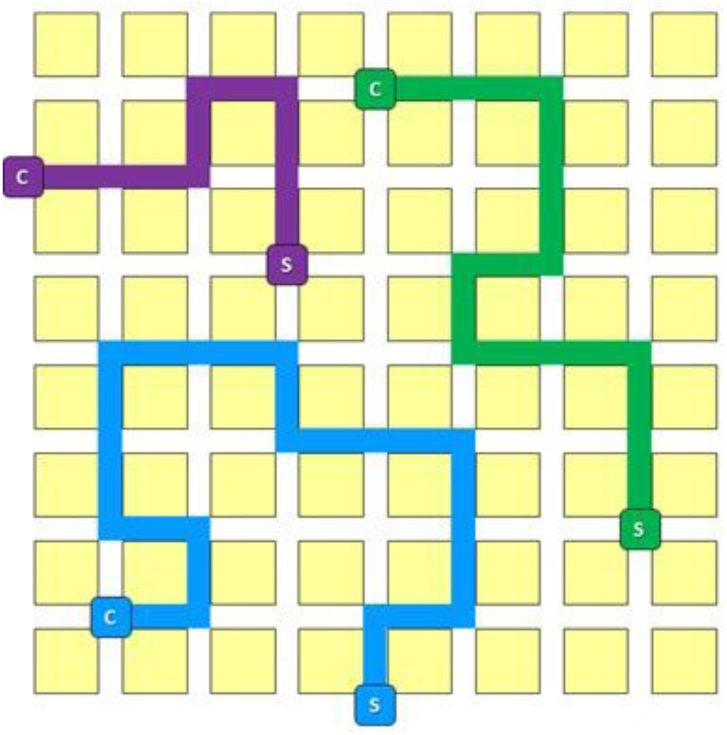
Depiction of three paths in our spatial memory tasks.

**Figure 2.**
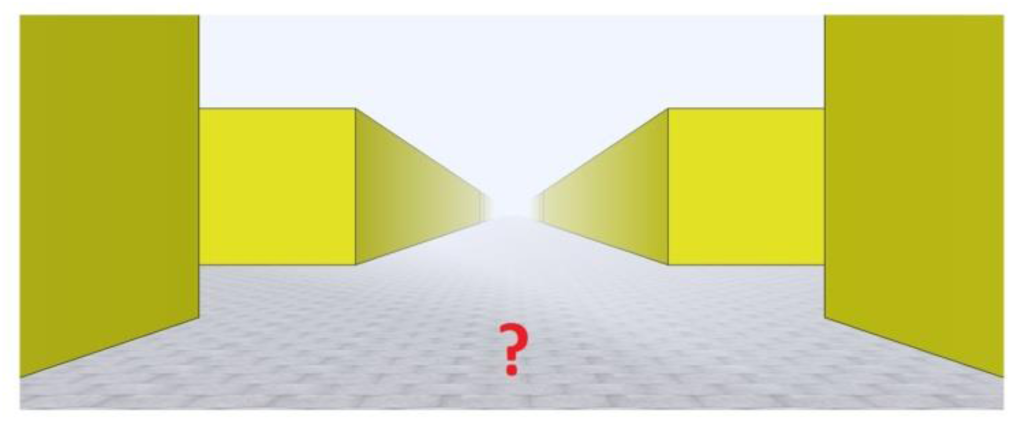
3D spatial memory task computerized environment.

### Procedure

Considering type of stimulation, our experiment had two experimental conditions repeated across all participants in the counterbalanced order. For every participant there was a time gap of at least 7 days between the 7-day follow-up test following the first session and the next session, so every tDCS session had the status of a single stimulation session. In one session participants received real while in the other they received sham stimulation. To assess immediate effects of tDCS treatment, we administered to participants 2D and 3D spatial memory tasks in duration of 15 minutes right after the stimulation. To assess potential long-lasting effects participants performed the same spatial memory task retest 24 hours and 7 days after the stimulation. Learning block was omitted in retest versions of the spatial task. Participants had a quick opportunity to remind the paths before each retest, when all three paths were showed together on the screen for three seconds.

## Results

### Immediate tDCS effects

Average (both conditions together) raw score of 2D recall immediately after the stimulation was 22.61 (75.4% accuracy), while average raw score of 3D task was 15.30 (51% accuracy). Descriptive statistics of spatial memory performance for each of the experimental conditions are presented in Table 1. Results of 2×2 repeated measures ANOVA (anode / sham; 2D / 3D) showed that 3D recall task was significantly harder (F(1, 21) = 198.547, p = .00) than 2D recall task. However, there was no significant effect of tDCS (F(1,21) = 2.08, p = .164), nor the significant effect of the task – tDCS interaction (F(1,21) = 0.152, p = .701).

**Table 1.**
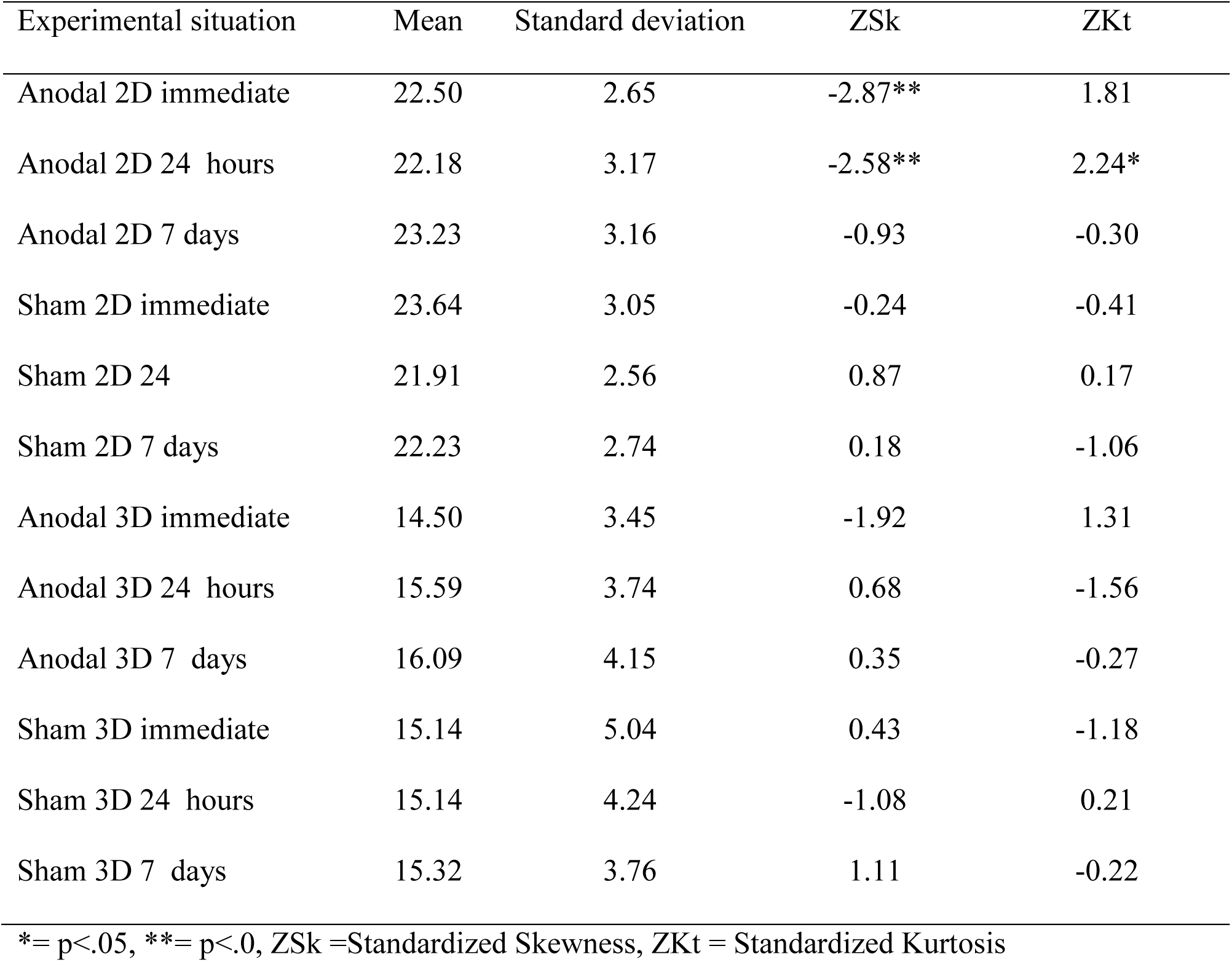
Spatial memory performance for all experimental conditions

### Long-term tDCS effects

In order to assess possible long-term effects of tDCS we conducted 2×3 repeated measures ANOVA (anode / sham ; immediate / 24 hours / 7 days) on 2D and 3D scores separately. No significant effects of tDCS were obtained on either 2D recall task (F(1,21) = 0.012, p = .914) or 3D recall (F(1,21) = .159, p = .694). In spite of slight trend of the average data to show an improvement of the results after 7 days in both 2D and 3D scores following tDCS (while they remained more or less the same in all three measurement following sham), the effect of time did not reach significance level for either 2D task (F(2,42) = 2.945, p = .064) or for 3D task (F(2,42) =1.202, p = .311) Finally, there was no significant effects of interactions: F(2,42) = 2.595, p = .087 ; F(2,42) = 0.780, p = .465 for 2D and 3D tasks respectively.

## Discussion

In this experiment we examined whether tDCS over right PPC can enhance spatial memory performance in two-dimensional and three-dimensional computerized environment. Additionally, besides immediate effects, we investigated possible long-term effects of tDCS after 24 hours and 7 days. No significant tDCS effects on spatial memory were observed in any of the three time points. This was the case with both 2D and 3D spatial memory recall measures.

Even though tDCS is a promising technique for drawing causal conclusions about the role of distinct brain regions in different cognitive functions there are studies challenging its effects. A recent meta-analysis (Horvath, Forte, & Carter, 2015) suggested that there was no convincing evidence for reliable effects of single session tDCS on cognitive functions in healthy adults. Still, there are several studies reporting significant cognitive enhancement by tDCS (Brunoni & Vanderhasselt, 2014; Hill, Fitzgerald, & Hoy, 2016; Summers et al., 2016). Results of our study do not provide evidence in favor of single session tDCS inducing relevant cortical functional change to affect performance on spatial memory.

On the other hand, there may be other reasons for the lack of tDCS effects in our study. There is consistent evidence about aging related decline in spatial memory functions, especially in allocentric framing. This was confirmed by results of meta-study (Colombo et al., 2017) that showed consistent decline of allocentric framing in healthy older adults. Potential explanation for this phenomenon is attenuated hippocampal activation, both at encoding and retrieval (Antonova et al., 2009). Thus, it is possible that we did not have appropriate sample for taping this effects. It would be reasonable to repeat this study on a sample of older participants. Another reason of possible attenuation of the effects in this study is complexity of spatial memory task that has been used. Namely, if performance on that task has been dependent on multiple cognitive processes, it is likely that electrode montage over P4 did not affect all neural structures that have substantial role in performance on our task. Finally, it should be considered that there is stronger empirical evidence about hippocampal role in spatial memory then it is for PPC, and that, due to its subcortical anatomical position, hippocampus is practically inaccessible for direct stimulation by non-invasive brain stimulation methods. Nevertheless, there is evidence that noninvasive stimulation of PPC can affect excitability changes in hippocampus by activating cortical-hippocampal brain network (Wang et al., 2014; Wang & Voss, 2015). Thus, it is not clear whether the lack of effects obtained in this study were due to not-strong-enough stimulation of hippocampus or due to the complexity of measure derived from spatial memory tasks that we had used, or combined effect of the both. Finally, in the literature that has been published so far not only that there is lack of empirical evidence about effects of non-invasive brain stimulation techniques on human spatial memory, but also there are not many consistent studies about tDCS effects over P4 on cognitive performance in general. Therefore, at present, it is hard to discuss further the meaning of our results. Some of the previous findings reported lack of positive effects of single session anodal tDCS over P4 on working memory (Berryhill, Wencil, Coslett, & Olson, 2010). On the other hand, there is evidence (Bjekić, Čolić, Živanović, Milanović, & Filipović, 2019) for memory performance enhancement by tDCS application over P4. Within this empirical background, our results remain unclear for now; there is certainly a need to reconsider our spatial memory measures and possible different electrode montage in further studies.

## Conclusion

To conclude, this study revealed no significant effects of single session tDCS over right PPC on spatial memory performance. Also, we did not observe any significant long-lasting effects (after 24 hours and 7 days) of a single tDCS treatment to spatial memory performance. The plausible explanation of the results we obtained stays unclear. Future studies should focus on using another multiple spatial memory measures and consider alternative electrode montages.

